# The effect of functionally-guided-connectivity-based rTMS on amygdala activation

**DOI:** 10.1101/2020.10.13.338483

**Authors:** L. Beynel, E. Campbell, M. Naclerio, J.T. Galla, A. Ghosal, A.M Michael, N.A. Kimbrel, S.W. Davis, L.G. Appelbaum

## Abstract

Repetitive transcranial magnetic stimulation (rTMS) has fundamentally transformed how we treat psychiatric disorders, but is still in need of innovation to optimally correct dysregulation that occurs throughout the fronto-limbic network. rTMS is often applied over the prefrontal cortex, a central node in this network, but less attention is given to subcortical areas because they lie at depths beyond the electric field penetration of rTMS. Recent studies have demonstrated that the effectiveness of rTMS is dependent on the functional connectivity between deep subcortical areas and superficial targets, indicating that leveraging such connectivity may improve dosing approaches for rTMS interventions. The current preliminary study, therefore, sought to test whether task-related, fMRI-connectivity-based rTMS could be used to modulate amygdala activation through its connectivity with the medial prefrontal cortex (mPFC). For this purpose, fMRI was collected on participants to identify a node in the mPFC that showed the strongest negative connectivity with right amygdala, as defined by psychophysiological interaction analysis. To promote long-lasting Hebbian-like effects, and potentially stronger modulation, 5Hz rTMS was then applied to this target as participants viewed frightening video-clips that engaged the fronto-limbic network. Post-rTMS fMRI results revealed promising increases in both the left mPFC and right amygdala, for active rTMS compared to sham. While these modulatory findings are promising, they differ from the a priori expectation that excitatory 5Hz rTMS over a negatively connected node would reduce amygdala activity. As such, further research is needed to better understand how connectivity influences TMS effects on distal structures, and to leverage this information to improve therapeutic applications.

## 1. Introduction

Repetitive transcranial magnetic stimulation (rTMS) is a non-invasive brain stimulation approach that uses rapidly changing magnetic fields to modulate neuronal activity underneath a stimulating coil. rTMS is approved by the U.S. Food and Drug Administration as a therapy for treatment-resistant depression and obsessivecompulsive disorders, and has also been proposed as a potential treatment for patients with posttraumatic stress disorders (PTSD). As reported in a recent comprehensive review published in a book chapter (Beynel, Appelbaum, & Kimbrel, 2020) 13 studies have been published that attempt to test rTMS effects on PTSD symptoms and/or pathophysiology. Collectively, these studies demonstrate significant, but modest PTSD symptom improvements from rTMS treatment. Surprisingly, however, across this literature varying the rTMS protocols, such as the use of inhibitory or excitatory pulse sequences or stimulation to the right or left cortical hemispheres have generally been found to produce equivalent effects on PTSD symptom improvement. One possible explanation for this consistent PTSD symptom improvement irrespective to rTMS administration protocol may be attributed to the fact that the effects of rTMS on brain circuitry may propagate across multiple brain regions involved in PTSD pathophysiology. Indeed, while rTMS effects are often assumed to be quite focal and superficial, with a depth penetration to about 2 cm below the scalp (Deng, Lisanby, & Peterchev, 2013), recent neuroimaging studies demonstrate modulation of interconnected brain regions (Bestmann, Baudewig, Siebner, Rothwell, & Frahm, 2003), including deep brain structures, which lie beyond the spread of the magnetic field (Vink et al., 2018). In a systematic review of 33 studies with baseline and post-rTMS measures of fMRI resting-state functional connectivity, it has been found that rTMS can induce significant changes in brain connectivity that spread both within and between functional brain networks (Beynel, Powers, & Appelbaum, 2020)

In light of these observed effects on BOLD signal and functional connectivity, recent studies are attempting to indirectly target distal brain areas through their resting-state functional connections with accessible, proximal cortical areas. This approach has been used successfully to modulate hippocampus (Wang et al., 2014), insula (Addicott et al., 2019; Li et al., 2017) and amygdala (Riedel et al., 2019). Such studies have demonstrated that “connectivity-based” rTMS may provide a promising approach to modulate deep brain regions, which is highly relevant when using rTMS as a treatment for psychiatric disorders that stem from fronto-limbic dysfunction and necessitate ways of modulating affected deep brain structures.

In this study, we build upon the current state-of-the-art to implement task-related functional connectivity-based rTMS. For this purpose, we derive individualized rTMS targets through application of psychophysiological interaction (PPI) analysis, in order to determine a location in the medial prefrontal cortex (mPFC) that shows maximal negative functional connectivity with the right amygdala, as participants passively viewed frightening or neutral pictures. Using this target, 5 Hz rTMS was applied ‘online’ as participants watched frightening video clips alternating with resting periods to engage the fronto-limbic network and induce Hebbian-like plasticity (Luber & Lisanby, 2014), under the hypothesis that such excitatory stimulation to the negatively connected node would strengthen the negative connectivity between the mPFC and the amygdala and consequently inhibit amygdala activity. It was also expected that, through Hebbian-like plasticity mechanisms, subjects experiencing the strongest feelings of fear, as reflected by changes in heart rate collected during the video-clips, would be the ones who show the strongest rTMS-induced changes in amygdala activation.

## 2. Methods

### 2.1. Participants

Fifty-two healthy young adults (18-35 years old) were contacted to participate in this this single blind, randomized placebo-controlled, two-visit study. The study was pre-registered on ClinicalTrials.gov (NCT03746405), and approved by the Duke University Health System Institutional Review Board (#Pro00101172). After the first phone screen, 27 participants declined to participate, and twenty-five were included in the study. During the first visit, participants completed an eligibility screening to ensure they did not have any contraindication to TMS (TASS, Keel, Smith, & Wassermann, 2001) or to MRI, followed by a psychiatric screening using the Mini-International Neuropsychiatric Interview (Sheehan et al., 1998) to ensure they did not have any disqualifying psychiatric disorders. They were then asked for a urine sample to ensure that the participants were not under the influence of any substances that could lower their seizure threshold and were not pregnant. Participants who made it through the above screenings (n = 23) were included in the study. Subsequently, three participants withdrew from the study due to scheduling conflicts, resulting in 20 participants completing the first visit (see **Figure 1** for consort diagram).

**Figure 1:**
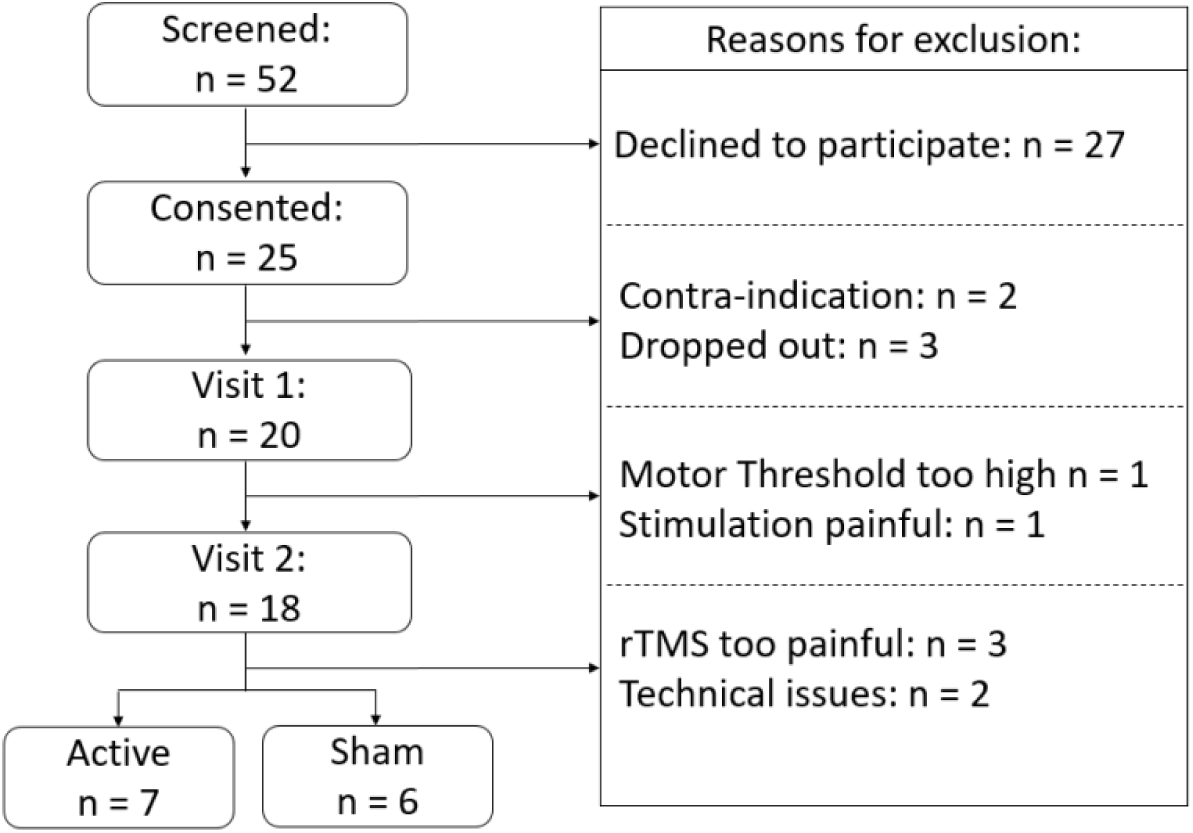
Consort diagram showing the recruitment, exclusion and inclusion numbers.

### 2.2 Experimental procedure

This study consisted of two experimental visits (**Figure 2A**). On the first visit, following screening and consenting, resting motor threshold (rMT) was determined and an MRI session was conducted to locate each participants rTMS target. During the second visit, which occurred no more than a week after the first visit, online rTMS was applied to individualized target locations as subjects viewed emotionally arousing video clips, immediately followed by a second MRI acquisition. rTMS effects were assessed by comparing fMRI BOLD signal and functional connectivity changes between these two visits. The following sections give greater details about each component of this procedure.

**Figure 2:**
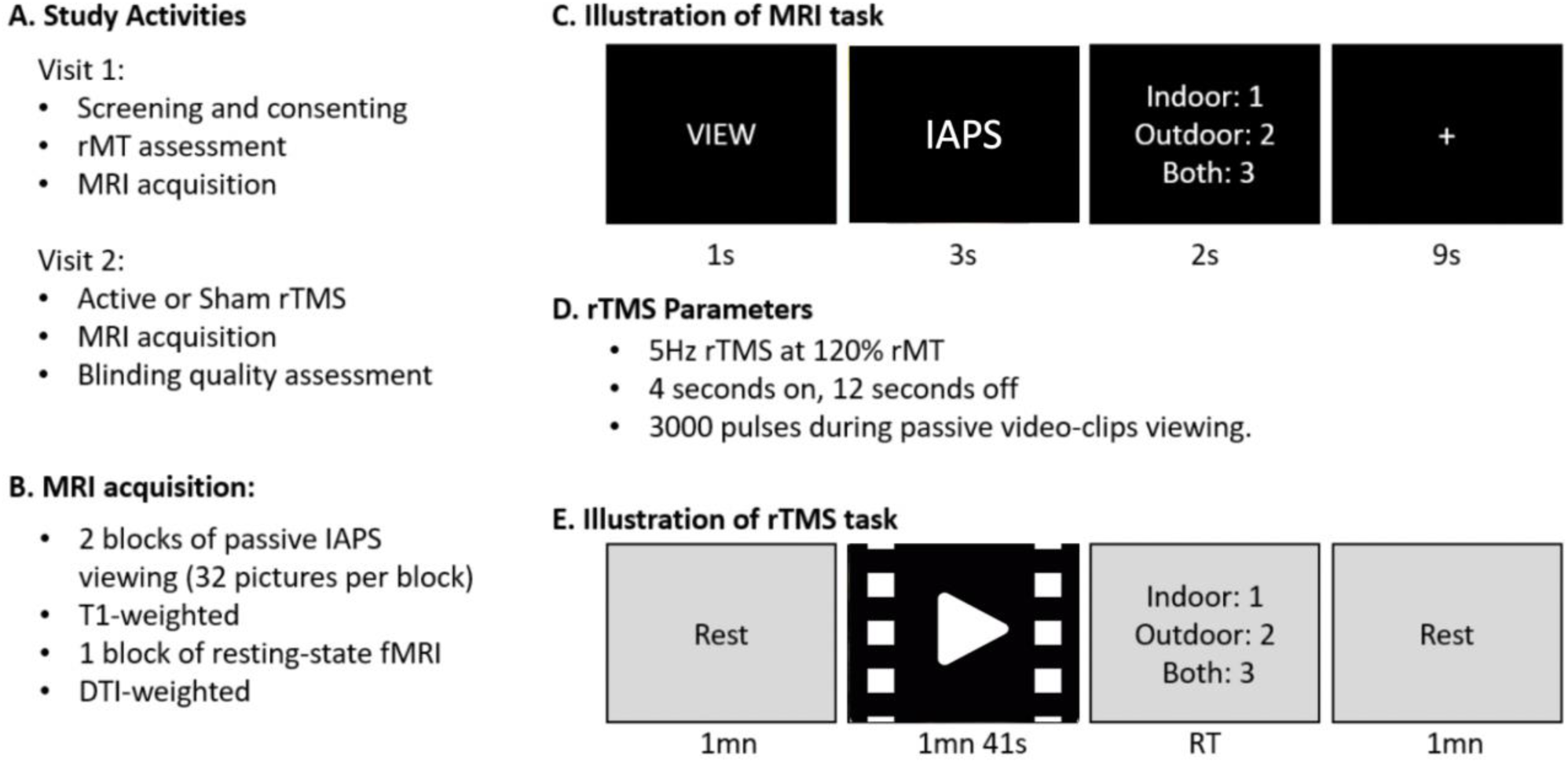
A) Activities done during each study visit. B) Scans during MRI acquisition. C) Illustration of MRI task showing passive viewing of IAPS pictures. D) rTMS parameters. E) Illustration of rTMS task showing passive viewing of frightening video-clips from the Schaefer et al. database. In order to maintain attention in both the MRI and rTMS tasks, participants were asked to rate whether the scene was indoor, outdoor, or both.

#### 2.2.1. Resting Motor Threshold (rMT)

rMT was performed with an active/placebo figure-8 coil (A/P Cool-B65) and a MagPro X100 stimulator with MagOption (MagVenture, Denmark), while the coil position was continually monitored through a stereotaxic neuronavigation system (Brainsight, Rogue Research, Canada). To define rMT, electrodes (Neuroline 720, Ambu) were placed on the right first dorsal interosseous (FDI) muscle in a belly-tendon montage and motor evoked potentials (MEPs) were recorded through the neuronavigation system. The motor ‘hot spot’ was defined as the position over the left motor cortex that elicited the greatest MEP in the right FDI. rMT was then defined as the TMS pulse intensity producing 50μV peak-to-peak MEP amplitude, using a maximum likelihood estimator (TMS Motor Threshold Assessment Tool, MTAT 2.0, http://www.clinicalresearcher.org/software.htm). Of the twenty participants who underwent rMT assessment, one was excluded because reported pain and a second was excluded because their rMT was above 83% of maximum stimulator output (MSO) and therefore at 120% of rMT would have exceeded the possible device output during the rTMS session. The remaining eighteen participants were then randomized into either active or sham rTMS group, and contacted to participate in the second visit.

#### 2.2.2. MRI acquisition

Participants completed a 30-minute MRI session (**Figure 2B**) that included a T1-weighted anatomical image (3D-T1-weighted echo-planar sequence, acquisition matrix = 256 mm^2^, time repetition [TR] = 7148 ms, time echo [TE] = 2.7 ms, field of view [FOV] = 256 mm^2^, spacing between slices = 1 mm, 196 slices), and a diffusion tensor imaging scan (acquisition matrix = 256mm^2^, TR = 17000 ms, TE = 91.4 ms, FOV = 256 mm2, spacing between slices = 2 mm, b-value = 2000 s/mm^2^, diffusion-sensitizing directions =25). Three runs of coplanar EPI functional images were also acquired with an oblique axial orientation including a resting-state scan and two blocks of passive viewing of IAPS pictures using the same acquisition parameters: (acquisition matrix = 128 mm^2^, TR = 2000 ms, TE = 25 ms, spacing between slices = 4mm, 240 volumes = 8 minutes per block). MRI acquisitions were collected on the first visit and on the second visit after rTMS intervention.

Each fMRI task block consisted in 32 trials. On each trial, a ‘VIEW’ instruction was displayed on the screen for one second, followed by three seconds of a picture from the International Affective Picture System (IAPS, Lang & Bradley, 2007). Three types of emotional pictures were used: fearful (50% of the trials), neutral (40%), and happy (10%). The images that were selected have been found to effectively evoke the desired affective responses (Barke, Stahl, & Kröner-Herwig, 2012; Schneider, Veenstra, van Harreveld, Schwarz, & Koole, 2016). The ‘happy’ category was added to reactivate the fronto-limbic network and limit habituation. In order to ensure that participants paid attention to the stimuli, they were asked to perform, in less than 2 seconds, a shallow scene judgment by indicating whether the image presented was outdoor, indoor or both, by pressing 1, 2 or 3 on the button box (see **Figure 2C**). Finally, a fixation cross was presented for nine seconds to allow the hemodynamic response to return to baseline, before the start of the next trial. To have a shorter delay between the end of rTMS and our task of interest, the functional task was always performed first and followed by the anatomical scan. During the restingstate acquisition, participants were asked to keep their eyes opened and to look at a white fixation cross on a black background. This setup has been shown to produce high test-retest reliability (Patriat et al., 2013). During the functional scan, visual stimuli were back projected onto a screen located at the foot of the MRI bed using an LCD projector. Subjects viewed the screen via a mirror system located in the head coil and the start of each run was electronically synchronized with the MRI acquisition computer. Behavioral responses were recorded with a 4-key fiber-optic response box (Resonance Technology, Inc.). Scanner noise was reduced with ear plugs, and head motion was minimized with foam pads. When necessary, vision was corrected using MRI-compatible lenses that matched the distance prescription used by the participant

#### 2.2.3. MRI processing for targeting approach

Following the first visit, functional connectivity between the right amygdala and medial prefrontal cortex (mPFC) was determined using psychophysiological interaction (PPI) analysis. To do so, functional images were skull stripped, reoriented and corrected for slice acquisition timing, motion, and linear trend using the FMRIB Software Library (FSL, https://fsl.fmrib.ox.ac.uk/fsl/fslwiki). Motion correction was performed using FSL’s MCFLIRT, and six motion parameters were then regressed out of each functional voxel using standard linear regression. Images were then temporally smoothed with a high-pass filter using a 190s cut off and normalized to the Montreal Neurological Institute (MNI) stereotaxic space.

Separate events were modeled for the viewing of the instructions (duration: 1s), rating (duration: 2s), and each of the emotional picture categories (fear, happy, neutral, duration: 3s), all with an onset at the beginning of the event, as recorded by the Matlab script used to launch the MRI acquisition. At the first level, functional data were analyzed as individual runs, using a general linear model (GLM) in which trial events were convolved with a double-gamma hemodynamic response function. The Fear > Neutral contrast was generated, allowing the identification of individualized statistical maps showing stronger BOLD activity when participants were seeing the fearful compared to the neutral pictures.

PPI analysis was then performed, for each functional run, following the FSL-PPI pipeline (https://fsl.fmrib.ox.ac.uk/fsl/fslwiki/PPIHowToRun). The design file from the previous BOLD analysis was used to generate the task regressor. To extract the time course of the right amygdala, the ‘fslmeants’ command was run by using the filtered functional data as the input and the right amygdala mask as defined by the AAL16 atlas (https://www.pmod.com/files/download/v35/doc/pneuro/6750.htm, ROI#42). These two events were then loaded into FSL: the task regressor as the psychological regressor, and the time course of the amygdala as the physiological regressor. The third event, the PPI was generated as the interaction between the task regressor and the amygdala. The remaining task regressors (happy, view and rate) from the original BOLD analysis were also included. The second level analysis was then performed to collapse information from both runs using a fixed-effects model. The subsequent statistical map was then moved back from MNI space to individual space using a linear registration (FLIRT). This map was then overlaid on the anatomical image on the neuronavigation software, and the region within the mPFC mask showing the strongest negative z-value was defined as the TMS target.

#### 2.2.4. Online rTMS procedure

During the second session, 5Hz rTMS was applied at 120% rMT over the mPFC target location, with the TMS coil handle was pointing upward. Stimulation was applied for 40 minutes, in trains of 4 seconds separated by intertrain intervals of 12 seconds (**Figure 2D**). These parameters replicate the ones used by Philip et al. (2018) who demonstrated significant connectivity changes between amygdala and mPFC in patients with posttraumatic stress disorders. Each participant received either active or somatosensory-matched sham stimulation, with the random allocation and assignment defined after each participant’s first visit. Sham stimulation was applied using the same coil in placebo mode, which produced similar clicking sounds and somatosensory sensation (via electrical stimulation with scalp electrodes) as the active mode, but with a greatly attenuated magnetic field that was shielded from the skull. To increase tolerability, lidocaine cream was applied on participant’s forehead before starting the experiment; and a ramp-up procedure was used by starting the stimulation at a very low intensity (10% MSO) and increasing it by 5% step during each rTMS inter train interval.

Given the importance of state-dependency on rTMS effect (Silvanto & Pascual-Leone, 2008) and in order to promote Hebbian-like plasticity, the stimulated fronto-limbic network was engaged during rTMS through the passive viewing of frightening movie clips chosen from a validated database and have been shown to reliably and effectively evoke feelings of fear (Schaefer, Nils, Sanchez, & Philippot, 2010). Each video-clip was followed by a shallow scene judgment (indoor, outdoor or both). Before the first movie clip and between the subsequent clips, there was a one-minute period where participants were instructed to rest (**Figure 2E**). Electrocardiography (ECG) was acquired throughout this task (LabChart, ADInstrument).

#### 2.2.5. MRI processing for group analysis to assess rTMS effect on amygdala activation and connectivity changes

Analysis of the rTMS effects on amygdala activation during passive viewing of IAPS pictures were done to emphasize emotional content of the images. As such, ‘VIEW’ events were not included, and the duration of the picture event was increased from 3 to 6 seconds, therefore including the rating event, and allowing a better representation of the hemodynamic response to implicit emotional processing of the image. The happy and neutral IAPS pictures were collapsed together and labeled as the “other” emotion which were contrasted with the fearful IAPS images. To assess connectivity changes, the ‘fear versus other” contrast was extracted from the BOLD analysis and used as the task regressor. The time course of the right amygdala was extracted as the physiological regressor. The third event, the PPI was generated as the interaction between the task and physiological regressor. No other events were entered into the analysis. For each of these outcomes, a 2*2 ANOVA was conducted with Timing (Visit 1 and Visit 2) as the within-subject factor and Stimulation (Active or Sham rTMS) as the between-subject factor.

#### 2.2.6. Blinding quality assessment

At the end of the second visit, to assess the quality of stimulation blinding, participants were asked to guess whether they received active or sham rTMS, and to rate their confidence in their guess on a scale from 0, indicating that they are not confident at all, to 100 indicating high confidence.

#### 2.2.7. ECG processing

To test whether participants expressed physiological fear responses while passively watching the video clips, heart rate beats per minutes (BPM) and heart rate variability (HRV) were extracted from the ECG data for each movie and rest periods. BPM and HRV for the nine movies and resting periods were averaged separately. An ANCOVA was then performed between the Stimulation (Active or Sham) and the Condition (Movies and Rest), with data from the first resting period, acquired before rTMS, used as the covariate to control for individual differences. According to the state-dependency assumption, individuals benefiting the most from rTMS should be the ones most engaged in the task, as indicated by their physiological responses. In this study, the changes in BPM and HRV between movies and resting periods were used as indicator of engagement in the task, and correlated with changes in amygdala activation between the two visits.

## 3. Results

### 3.1. Tolerability and blinding

Although rTMS intensity was calibrated according to rMT for each individual and a slow ramp-up procedure was used to acclimate participant to the sensation, stimulation was still too painful for three participants who withdrew from the study. As such, 13 subjects (7 females and 6 males) completed the full protocol and were included in the final analysis. These individuals had a mean age of 23.6 years (SD = 3.01), with seven participants randomly assigned to active group and six to the sham group.

To assess the quality of the blinding process a chi-square test of independence was performed. This test did not reveal any dependence between participants actual and guessed group assignments (p = 0.42). However, a significant relationship was found between the confidence of their guess, and the true delivered stimulation (p < 0.05). The numerical values (**Table 1**) indicate that participants tend to guess that they received active stimulation for both the active and sham true stimulation condition, which support a good blinding quality.

**Table 1:** Count of participants’ guess and averaged confidence level in their guess (from 0%: not confident to 100%: definitely confident in their guess) as a function of the true delivered stimulation.

|  |  | Participants' Guess |  | Confidence Level of Guess |  |
| --- | --- | --- | --- | --- | --- |
|  |  | Active | Placebo | Active | Placebo |
| Actual Stimulation | Active | 6 | 1 | 79.2 | 10.0 |
|  | Sham | 4 | 2 | 57.5 | 27.5 |

### 3.2. MRI and electrophysiological changes

#### 3.2.1. BOLD changes in Fearful versus Other contrast

Results from the ANOVA revealed a main effect of Timing with significantly less activations on Visit 2 than on Visit 1. In particular, bilateral amygdala (**Figure 3A**) showed a main effect of Stimulation, with stronger activation for Active than Sham rTMS in the prefrontal cortex (**Figure 3B**), and a significant interaction between Timing and Stimulation (**Figure 3C**).

**Figure 3.**
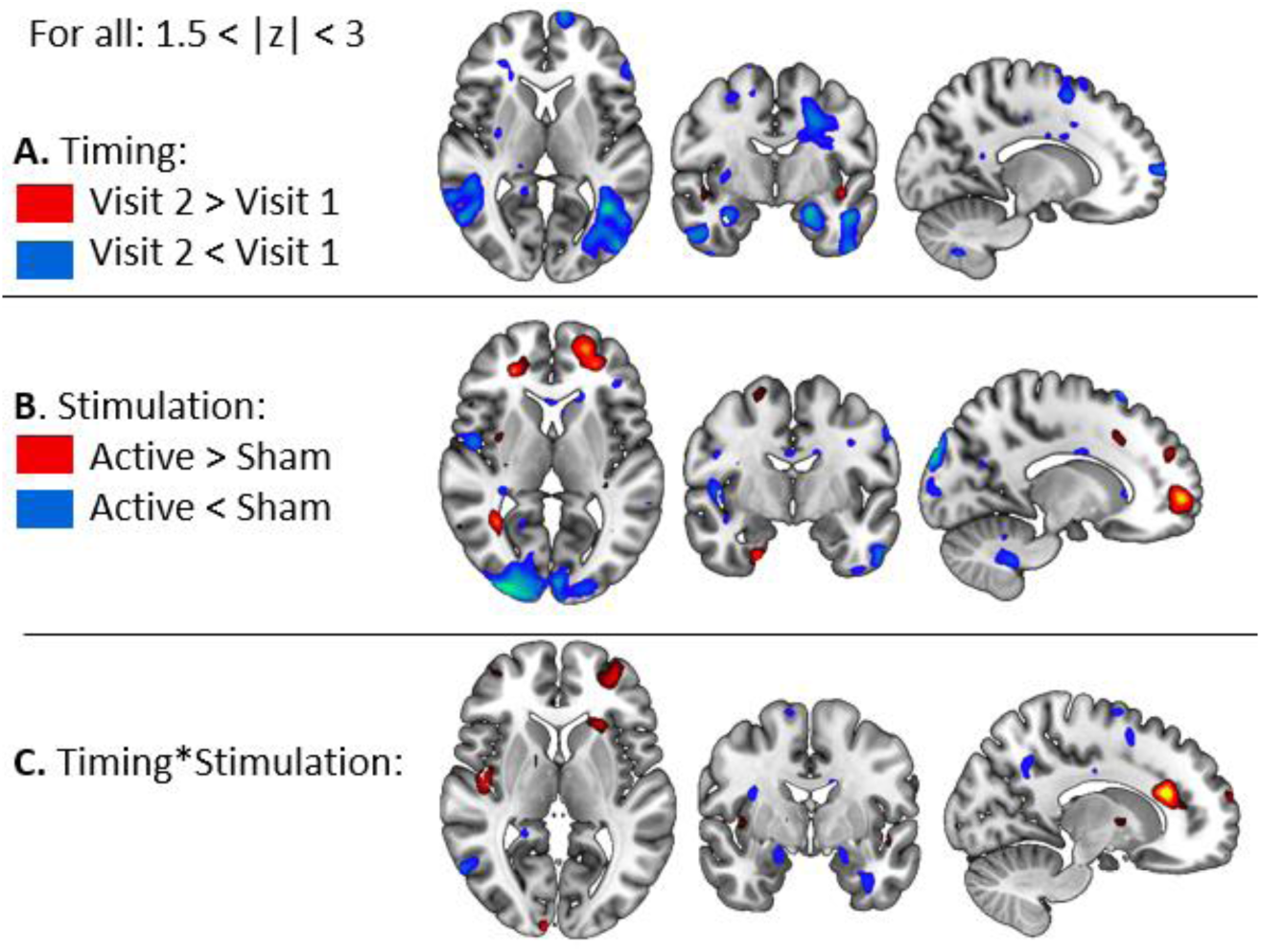
Results from the 2*2 ANOVA on BOLD signal. A) Main effect of Timing (Pre versus Post rTMS). B) Main effect of Stimulation (Active versus Sham rTMS), and C) Interaction between Timing and Stimulation.

Since significant differences were found in the main analysis, we then performed t-tests contrasts to better assess each difference. First, BOLD signal in the Fear versus Other contrast was computed in each visit separately. During the first visit, the presentation of frightening IAPS pictures increased amygdala activation when compared to other pictures (**Figure 4A**). This result demonstrated that the task induced expected affective brain changes and allowed for investigation of subsequent amygdala changes after rTMS. During the second visit, it was found that the amygdala was no longer significantly activated (**Figure 4B**), likely due to habituation, which has previously been reported in similar studies (Breiter et al., 1996), and likely explains the main effect of timing found in the larger analysis. The interaction was then decomposed to assess the influence of stimulation condition, during Visit 2. Given the limited number of subjects in this analysis (7 versus 6) the significance threshold level was decreased from 1.5 to 1. Here, it was found that when compared to subjects receiving electrical sham stimulation, subjects receiving active rTMS displayed stronger activation both in the stimulated left mPFC, and in the indirectly targeted right amygdala (**Figure 4C**, green shaded ROIs). This finding indicates that active rTMS, applied over the mPFC counteracts habituation and provides evidence that connectivity-based rTMS is able to modulate amygdala, in a manner that is specific to the hemisphere of stimulation.

**Figure 4:**
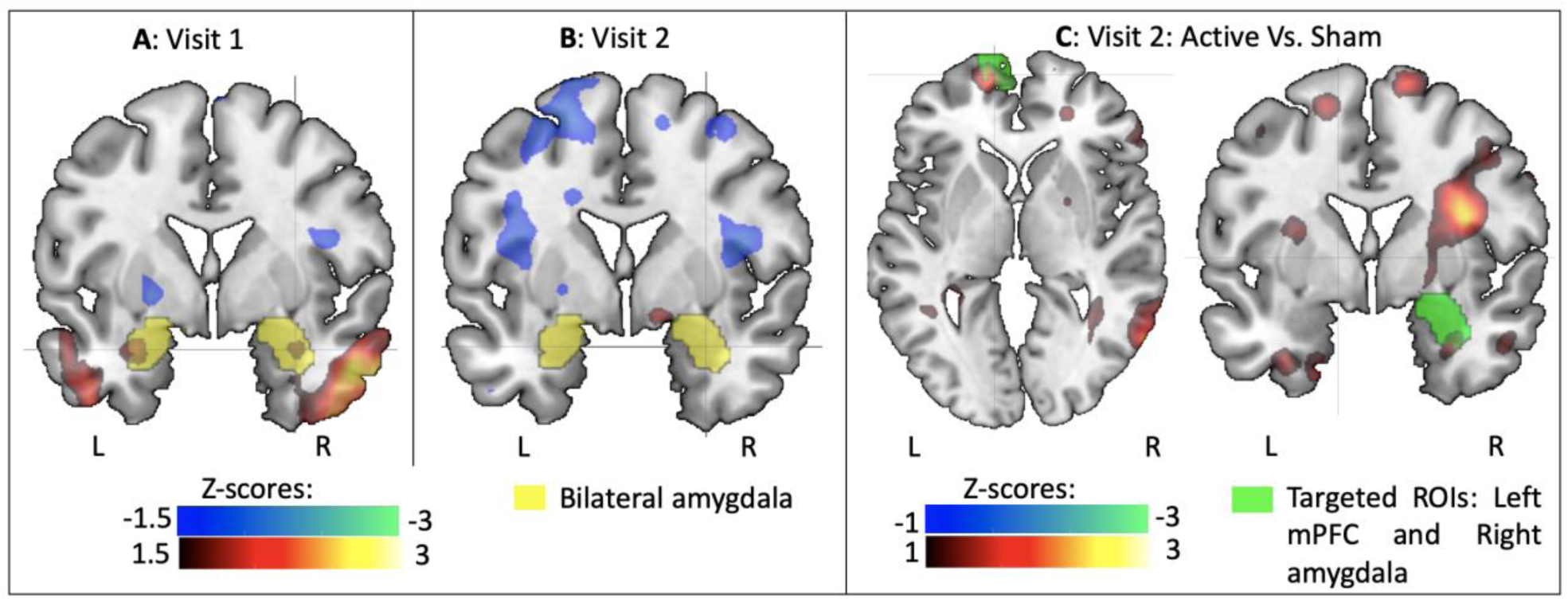
BOLD signal analysis with for the Fear versus Other contrast on Visit 1 (A), Visit 2 (B), and comparison of the effects of Active versus Sham rTMS on the second visit (C). Heat maps indicate Z-scores, blue colors indicate a decrease in BOLD signal and red colors indicate increase. Yellow shading represents the amygdala; and green shading represents masks used to define left mPFC and right amygdala.

To more specifically test the effect of rTMS on BOLD activity in the right amygdala, an ROI analysis was performed. The right amygdala ROI was first generated by combining the group activation from Visit 1 (z > 1.5) and the anatomical mask from the AAL atlas. Z-scores in the Fear vs. Other contrast within this ROI were then extracted for all subjects at each visit. Two independent t-tests were then performed on these z-scores to investigate the differences between the two groups. While, as expected, no differences were found on the first visit (p = 0.95), a significant difference was found on the second visit, with subjects in the active group showing significantly greater amygdala activation (mean = 0.67, standard deviation = 0.48) than subjects receiving sham rTMS (mean = −0.48, standard deviation = 0.87; p = 0.03) (**Figure 5**). This ROI analysis therefore confirmed the results from the whole-brain analysis with active rTMS significantly increasing activity in the right amygdala compared to sham rTMS.

**Figure 5:**
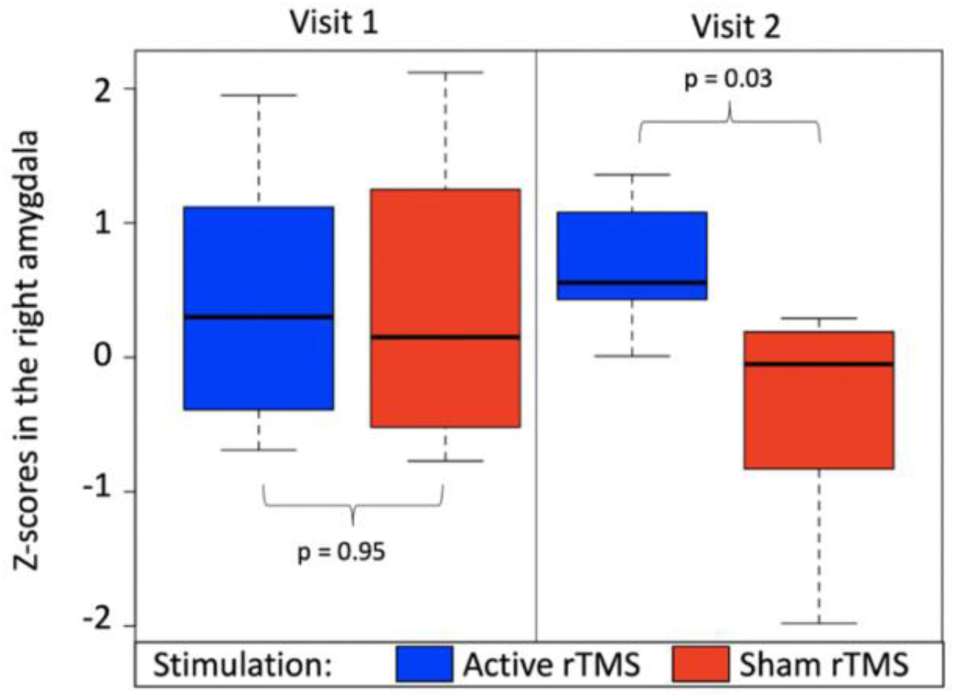
fMRI activation in the right amygdala (z-scores) obtained before rTMS (Visit 1) and after rTMS (Visit 2) for subjects receiving active stimulation (blue) or sham stimulation (red). The p-values are reported for independent t-tests comparing amygdala activation in each group, within each visit.

#### 3.2.2. Task-related functional connectivity change

Results from the ANOVA revealed a main effect of Timing with significantly stronger connectivity between the right amygdala and the whole brain on Visit 2 than on Visit 1 (**Figure 6A**). A main effect of Stimulation was also found with stronger connectivity after Active compared to Sham rTMS (**Figure 6B**) and a significant interaction between Timing and Stimulation (**Figure 6C**).

**Figure 6.**
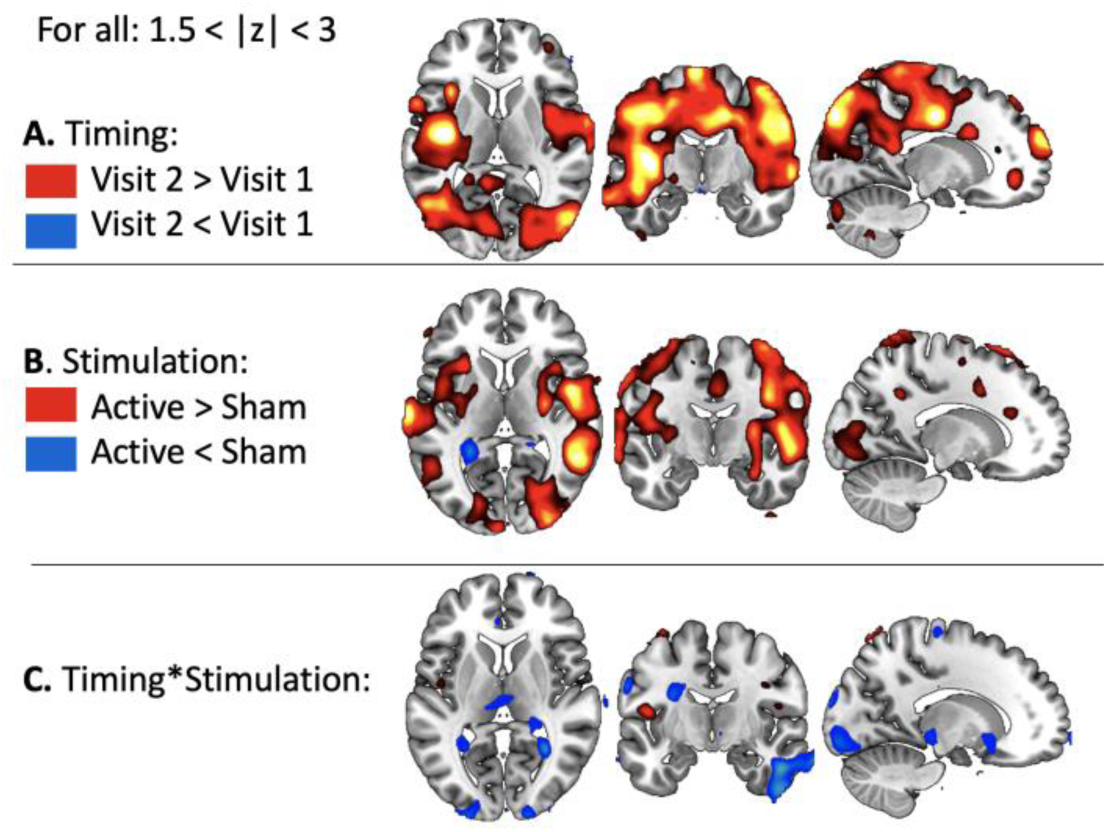
Results from the 2*2 ANOVA on task-related functional connectivity. A) Main effect of Timing (Pre versus Post rTMS). B) Main effect of Stimulation (Active versus Sham rTMS), and C) Interaction between Timing and Stimulation.

To better understand these effects, t-tests were conducted, and demonstrated that the connectivity pattern was reversed during the second visit, by switching from a negative connectivity between the right amygdala and the whole brain, during the first visit (**Figure 7A**) to a positive connectivity on the second visit (**Figure 5B**). When comparing the effects of Active and Sham rTMS in Visit 2, it was observed that these changes were due to increased functional connectivity with Active rTMS in the stimulated left mPFC (**Figure 7C**).

**Figure 7:**
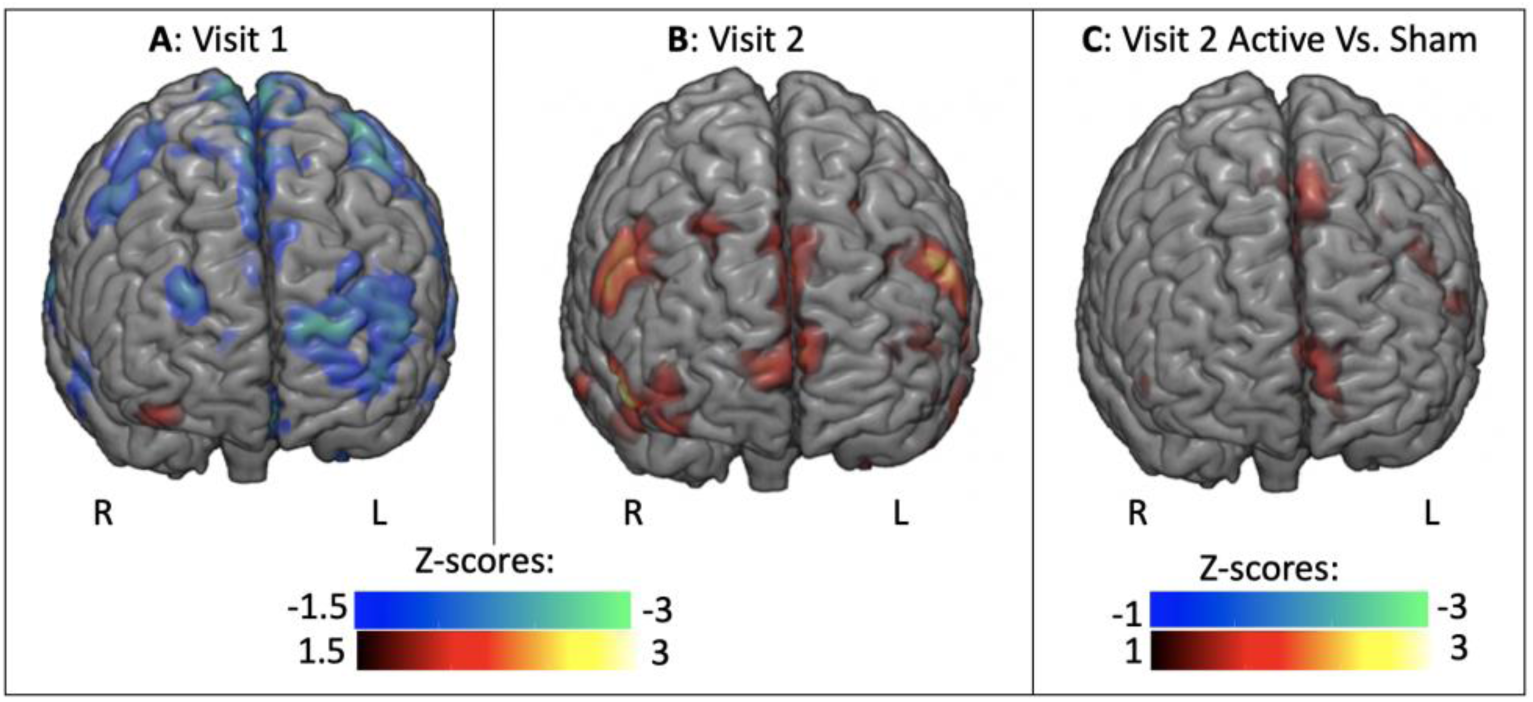
PPI analysis for the Fear versus Other contrast on Visit 1 (A), Visit 2 (B), and comparison of the effects of Active versus Sham rTMS on the second visit (C). Blue colors indicate decreased connectivity; and red colors indicate increased in functional connectivity.

#### 3.2.4. Beats per minutes and heart rate variability during movies and resting periods

Two mixed measures ANCOVAs were used to assess changes between frightening movies and rest conditions, and to test the effect of Stimulation on for BPM and HRV, using the first resting period as a covariate. Contrary to our assumption, no significant differences were found on the number of beats per minutes between the two conditions of interest (Movie: 67 ± 11.14 versus Rest: 66.28 ± 10.82, F(1,12) < 1). No differences were found between Stimulation (Active: 65.29 ± 12.32 versus Sham: 67.82 ± 9.52; F(1,12) < 1), and the interaction between these two factors was not significant (F(1,12) = 1.25, p = 0.29) (see **Table 2** for numerical values). The same pattern of null results were found for HRV with no effect of Condition (Movie: 0.93 ± 0.18 versus Rest: 0.95 ± 0.18, F(1,12) < 1) or Stimulation (Active: 0.95 ± 0.18 Sham: 0.92 ± 0.17), or interaction between Stimulation and Condition (F(1,12) = 1.27, p = 0.28) (see **Table 2**).

**Table 2:** Averaged beats per minutes (BPM) and heart rate variability (HRV) during movies and resting period for participants who received active or sham rTMS.

|  | Movies |  | Rest |  |
| --- | --- | --- | --- | --- |
|  | BPM | HRV | BPM | HRV |
| Active rTMS | $65.19 \pm 12.88$ | $0.95 \pm 0.19$ | $65.39 \pm 12.77$ | $0.95 \pm 0.19$ |
| Sham rTMS | $68.31 \pm 11.59$ | $0.90 \pm 0.18$ | $66.17 \pm 11.23$ | $0.94 \pm 0.17$ |

According to the state-dependency assumption, it was expected that participants in the active group who experienced higher physiological arousal as measured by differences in BPM and HRV between movies and resting periods would show greater changes in amygdala activation. While no significant correlations were found, which may be due the small sample size and the lack of significant difference between movies and rest condition, the results seem to indicate that this assumption is true. Indeed, a positive relationship was found between changes in HRV and changes in amygdala activation, suggesting that individual showing larger variability between movies and rest conditions are the ones showing the strongest changes in amygdala activation (r = 0.32, p = 0.48). However, this relationship was negative when using BPM (r = −0.19, p = 0.68) (**Figure 8**).

**Figure 8:**
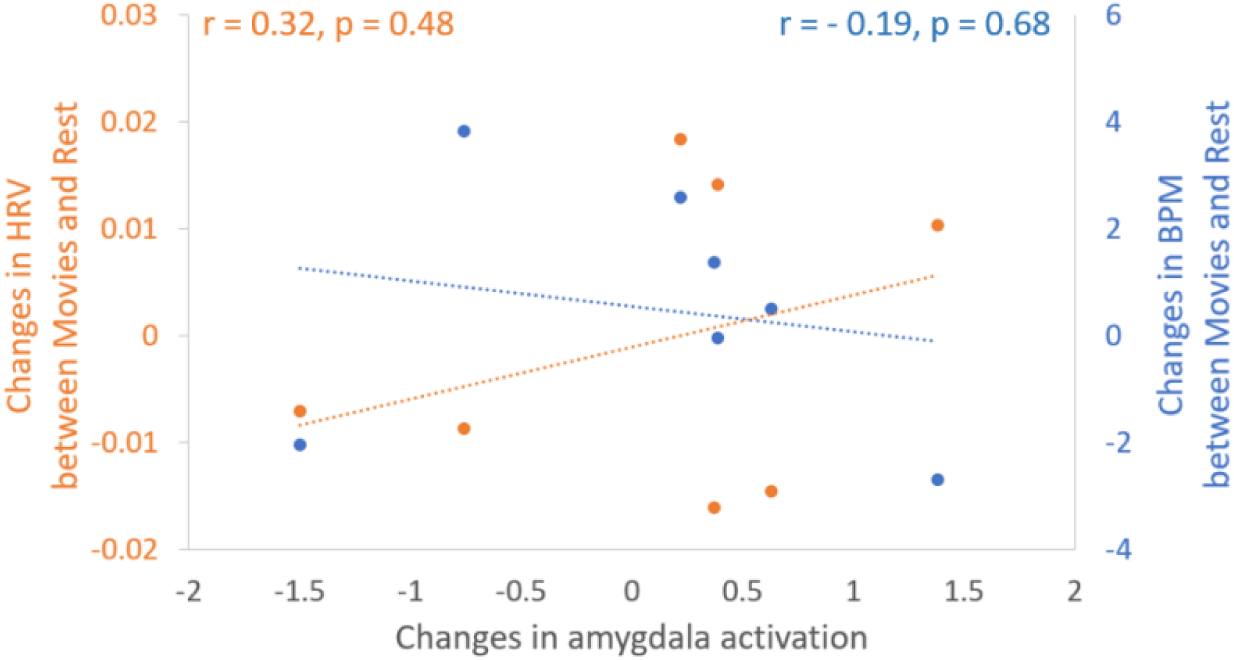
Scatterplot showing changes in amygdala activation between visit 2 (after rTMS) and visit 1 (before rTMS) on the x-axis, and changes between movies and resting periods for HRV (orange) and BPM (blue) on the y-axis.

## 4. Discussion

In this preliminary, proof-of-concept study, connectivity-based rTMS was applied over the node in the medial prefrontal cortex (mPFC) that showed the strongest negative connectivity with the right amygdala, as defined by a PPI analysis. It was anticipated that active rTMS over this target would reduce amygdala activity by strengthening the negative connectivity between these two regions. To improve rTMS efficacy, stimulation was applied online, while subjects were passively viewing frightening video clips from the Schaefer et al. database (Schaefer et al., 2010), separated by resting periods, under the assumption that engaging the fronto-limbic network while stimulating it would promote Hebbian-like plasticity.

The results offer a highly promising demonstration of the potential for connectivity-based target engagement that is feasible and practical. Interestingly, the results were opposite of what was expected, as (i) active rTMS did not strengthen negative connectivity between mPFC and amygdala, but instead reversed it, and (ii) the activity of the indirectly targeted right amygdala activity was increased, instead of decreased, after active relative to sham rTMS. However, as expected, changes in amygdala activity were found to be positively correlated with changes in heart rate variability, suggesting that the subjects benefiting the most from rTMS expressed the most physiological arousal and greater task engagement.

Understanding how the proximal rTMS effects propagate to distal structures remains highly complex and yet rarely explored in the literature. In a recent review article of 33 studies investigating rTMS effects on resting state functional connectivity (Beynel, Powers, et al., 2020), it was found that the common rTMS frequency-dependent heuristic observed with proximal brain structures was not typical of studies reporting downstream distal effects, with a majority of studies reporting increased functional connectivity after rTMS, independent of the stimulation frequency. Results from the current study point in this same direction. However, it is difficult to draw strong conclusions from our results, not only because the sample size was highly limited, but mainly because a large decrease in amygdala activation was observed between the two visits, probably due to task habituation (Breiter et al., 1996) which constitutes a large bias. Indeed, the targeting approach was based on the task-related connectivity between the activated amygdala and mPFC, but since the amygdala was not activated by this task during the second MRI acquisition, the mPFC does not need to exert his inhibitory control of the amygdala anymore. Therefore, it is impossible with these data to define whether the increase in amygdala activation observed between active and sham rTMS during the second visit was due to a connectivity-change or to an actual change in amygdala activity. Further research is needed to understand this link, by collecting fMRI right before and right after the rTMS intervention, and testing the interaction between stimulation frequency (low versus high frequency) and connectivity profile (stimulating a node positively or negative connected to the amygdala) on amygdala changes.

Regarding the electrophysiological data, while no differences were found on heart rate variability between the frightening video-clips and the interleaved resting periods, suggesting that participants did not experience high physiological arousal, results still seem to indicate that the subjects benefiting the most from rTMS were the ones showing the stronger changes between movies and rest. This result is in line with the state-dependency assumption (Silvanto & Pascual-Leone, 2008) and highlight the importance of controlling the subject’s state during rTMS to promote stronger efficacy. A potential way to improve this result for this specific study would be to add movies with positive valence that would prevent the chance of developing habituation effects.

To conclude, this study demonstrated the feasibility and promising effect of task-related connectivity-based rTMS on amygdala activation, and demonstrated the importance of state-dependency on rTMS efficacy. More research is required to reliably predict and leverage these distal effects for specific applications. If successful, these studies could pave the way to more powerful neurotherapeutic approaches that can be applied to patients with fronto-limbic cortical dysregulation, such as posttraumatic stress disorders.

## Acknowledgments

This research was funded by a Duke Institute for Brain Sciences Research Germinator Award

